# Targeting of the Mon1-Ccz1 Rab guanine nucleotide exchange factor to distinct organelles by a synergistic protein and lipid code

**DOI:** 10.1101/2022.08.14.503906

**Authors:** Eric Herrmann, Lars Langemeyer, Kathrin Auffarth, Christian Ungermann, Daniel Kümmel

## Abstract

Activation of the small GTPase Rab7 by its cognate guanine nucleotide exchange factor (GEF) Mon1-Ccz1 (MC1) is a key step in the maturation of endosomes and autophagosomes. This process is tightly regulated and subject to precise spatiotemporal control of MC1 localization. We here identify and characterize an amphipathic helix in Ccz1, which is required for the function of Mon-Ccz1 in autophagy, but not endosomal maturation. Furthermore, our data show that the interaction of the Ccz1 amphipathic helix with lipid packing defects, binding of Mon1 basic patches to positively charged lipids and association of MC1 with recruiter proteins collectively govern membrane recruitment of the complex in a synergistic and redundant manner. The data demonstrate that specific protein and lipid cues convey the differential targeting of MC1 to endosomes and autophagosomes. We reveal the molecular mechanism how MC1 is adapted to recognizes distinct target compartments by exploiting the unique biophysical properties of organelle membranes and thus provide a model how the complex is regulated and activated independently in different functional contexts.

## Introduction

Organization of intracellular processes in eukaryotes relies on the compartmentalization of biochemical processes to organelles and sub-organellar domains (1). Consequently, the targeting of proteins to the appropriate microcompartments underlies strict spatiotemporal control. Molecular on/off switches, namely reversible phosphorylation by kinases and phosphatases (2) or cycling of GTPases between GTP- and GDP-bound form (3), can regulate the activity and interactions of biomolecules. However, rather than control by single all-or-nothing signals, many systems utilize multiple cues and coincidence mechanisms for adaptive regulation (4, 5). The same functional modules can thus fulfill different functions in different settings. Unraveling the molecular basis that governs these processes is required for a mechanistic understanding of cellular functions.

The Mon1-Ccz1 (MC1) complex is a guanine nucleotide exchange factor (GEF) that activates the endosomal GTPase Rab7 (Ypt7 in yeast) (6–9). MC1-dependent activation of Rab7 is required for fusion of multi-vesicular bodies (MVB)/late endosomes and of autophagosomes to lysosomes (vacuole in yeast) (6, 10–13). Recently, MC1 was also implicated to support mitophagy (14). In yeast, the interactions of MC1 with Vps21 (a Rab5-like GTPase) (10, 15), Atg8 (the homolog of mammalian LC3) (11, 16) and phosphatidylinositol phosphate (PIP) lipids (13, 17, 18) were shown to be required for complex function. Homologous interactions were also described in mammalian cells. Likely, the differential association of MC1 to these localization cues contributes to the spatiotemporal control of its activity.

The structural characterization of MC1 provides a framework for a mechanistic analysis of its function. The complex belongs to the family of Tri Longin Domain (TLD) RabGEFs, which are characterized by a heterodimeric core complex (19, 20). The core subunits, Mon1 and Ccz1 in the case of MC1, are each composed of three conserved longin-type domain (LDs) that are arranged in a triangular fashion. Together they form a complex with pseudo two-fold symmetry (21) resulting in a two-layered architecture. The bottom layer, formed by the LD2s and LD3s, is required for proper subcellular localization of the complex. A large positively charged patch on the bottom layer mediates membrane binding, defining the orientation of MC1 on the membrane. The top layer, comprised of the LD1s, constitutes the catalytic center that promotes nucleotide exchange of Rab7/Ypt7 (22).

Our previous work addressed the GEF activity of MC1, revealing a unique lysine-insertion mechanism (22) and activation of the complex by its recruiter GTPases on model membranes (15). In this study, we focused on how MC1 membrane interactions and targeting to distinct cellular compartments are realized on a molecular level. We performed a detailed analysis of MC1 recruitment to membranes using *in vitro* reconstitutions and functional analysis *in vivo*. We identify binding to lipid packing defects as an additional targeting mechanism of MC1 that is essential specifically for its function in autophagy. Interestingly, different recruiter proteins and lipid binding cues act on MC1 in a synergistic, but also redundant manner, providing a mechanism how fine-tuning of activity and differential targeting to distinct organelles can be achieved.

## Results

### Mon1-Ccz1 binding to negative charges and packing defects in membranes

In previous work with the Mon1-Ccz1 complex from *Chaetomium thermophilum* (CtMC1), we demonstrated that CtMon1 LD2/3 are required for the recruitment of MC1 to PIP-containing liposomes and subsequent co-sedimentation (21). We noticed that CtMC1 also bound weakly, but reproducibly to neutral liposomes containing palmitoyl (16:0) - oleoyl (18:1) (PO) phospholipids (Fig S1A,B). This effect was more pronounced with liposomes that were composed of di-oleoyl (DO) phospholipids (Fig S1A,B). Using different truncation constructs of MC1 in liposome sedimentations, we found that LD2 and LD3 of CtCcz1 are required for the interaction of CtMC1 with neutral liposomes (Fig 1A,B). Because DO liposomes contain more unsaturated acyl chains they have more lipid packing defects than PO liposomes. We therefore concluded that CtMC1 contains a lipid packing sensing motif in the ‘localization layer’ of CtCcz1. In contrast, binding of CtMon1 with LD2/3 to liposomes containing 2% phosphatidylinositol-3-phosphate/1% phosphatidylinositol-3,5-bisphosphate (PIP) was independent of packing defects (Fig 1A,B). Thus, CtMon1 and CtCcz1 can both interact with membranes, recognize positive charges and lipid packing defects, respectively, and jointly mediate membrane binding of the complex and its recruitment onto lipid bilayers.

**Figure 1:**
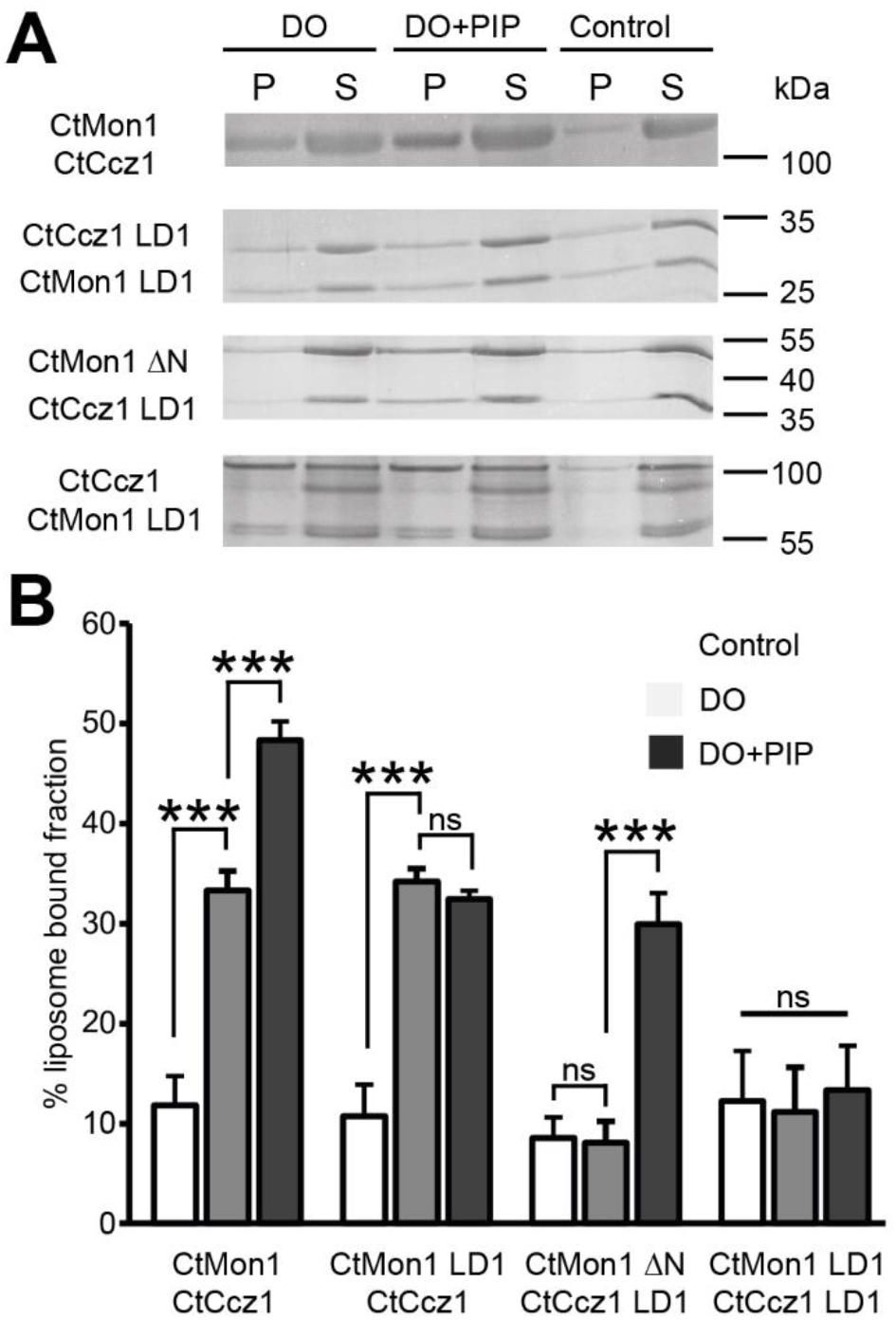
Lipid interactions of the MC1 complex. (**A**) Sedimentation assays of full-length CtMC1 and different deletion complexes with liposomes (400 nm) from a DO lipid mix with or without PIP (**B**) Quantification of A (n=4).

### Synergistic recruitment of MC1 by lipids and proteins

Next, we asked if the lipid binding properties of MC1 are conserved and therefore investigated the interaction of the Mon1-Ccz1 complex from *Saccharomyces cerevisiae* (ScMC1) with liposomes (Fig 2A,B). The yeast complex also showed robust binding to DO liposomes (high packing defects), but only weak interaction with PO liposomes (low packing defects). Furthermore, ScMC1 was efficiently recruited to PO+PIP liposomes (negative charges), recapitulating the findings obtained with CtMC1.

**Figure 2:**
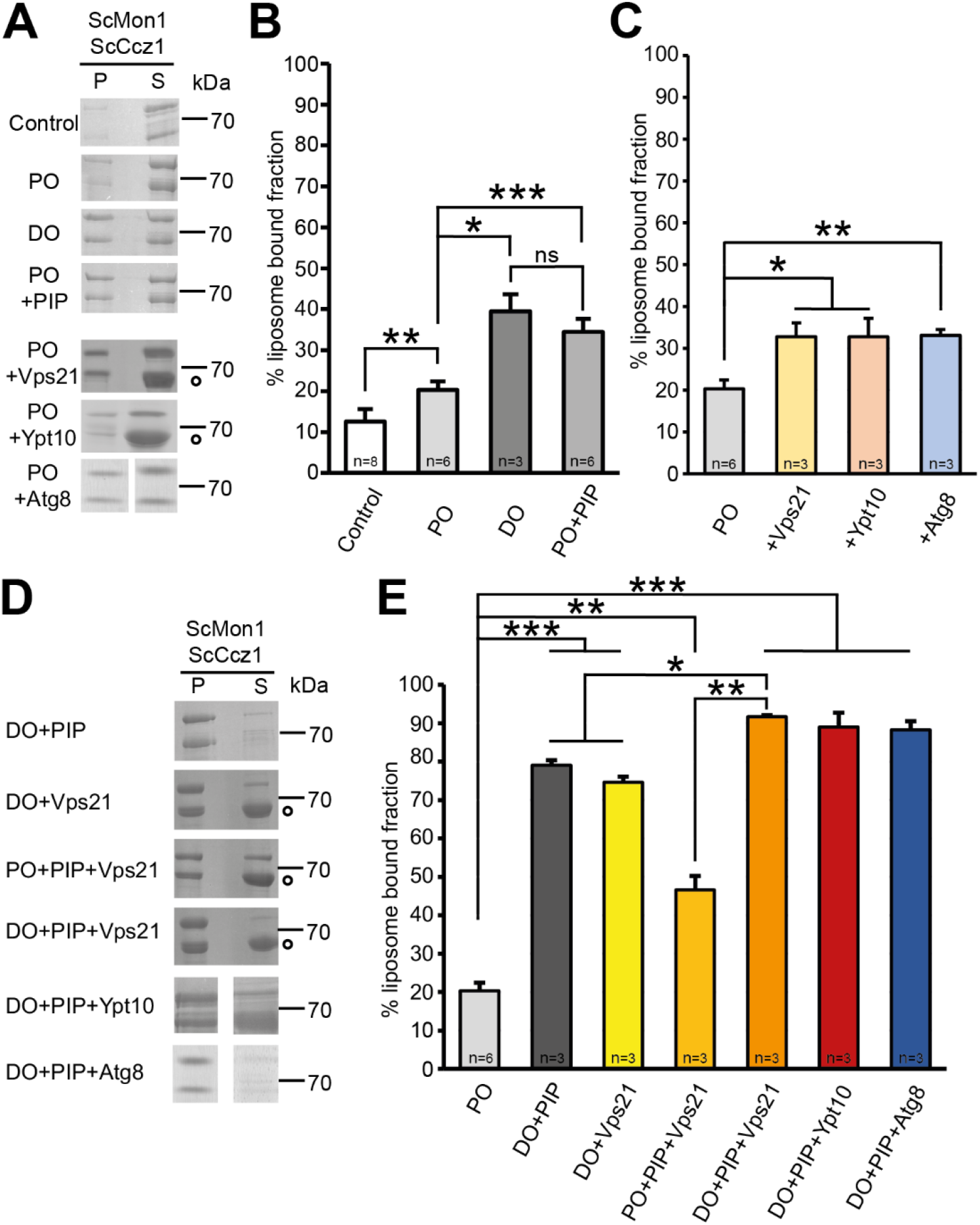
Triple targeting mechanism of the MC1 complex. (**A**) Binding of ScMC1 to liposomes from a PO, DO, and PO-PIP mix, and to PO liposomes with Vps21, Ypt10 or Atg8. The Mrs6 Rab escort protein from the prenylation reaction is marked by a circle. Sample without liposomes serves as control. (**B**) Quantification of ScMC1 binding to PO, DO, and PO-PIP lipososmes (n=3-8) (**C**) Quantification of ScMC1 binding to PO liposomes with Vps21, Ypt10 or Atg8 (n=3-6). (**D**) Binding of ScMC1 to PO or DO liposomes containing PIPs and recruiter proteins as indicated. (**E**) Quantification of G (n=3-6). Quantification data are presented as mean ±SD and the significance was calculated using Student’s t-test (* p<0.05, ** p<0.01, *** p<0.001, n.s. not significant).

ScMC1 was reported to require the interaction with recruiter proteins for its function, namely the autophagosomal marker Atg8 and the endosomal Rab5-like GTPases Vps21 and Ypt10 (11, 15). To test their influence on membrane targeting of ScMC1, we incorporated the naturally lipidated forms (prenylated Vps21 or Ypt10 and Atg8 coupled onto phosphatidylethanolamine (PE)) of the recruiter proteins into PO liposomes and measured binding in liposome sedimentation assays. Each recruiter promoted membrane binding to a similar extend, but relatively weakly (Fig 2A,C).

Neither recruitment cue alone - negative charges, lipid packing defects or a recruiter protein - was able to efficiently target ScMC1 to liposomes. We thus tested different combinations of liposomes with packing defects (DO lipids), negative charges (PIPs) and the recruiter GTPase Vps21. Two recruitment cues remarkably enhanced ScMC1 membrane binding, in particular when liposomes contained packing defects (Fig 2D,E, S1C). In particular, the binding observed with DO+PIP liposomes (high packing defects, negative charges) was much stronger, indicative of two independent lipid interaction sites that act synergistically. Membrane binding was further increased when all three recruitment cues were present on the liposomes (Fig 2D,E). The triple targeting set-up also led to almost complete ScMC1 membrane binding when Ypt10 or Atg8 were used as recruiter proteins.

### Role of PIP binding in vivo

To gain insight into the relevance of PIP binding in physiology, we sought to identify mutations that specifically abolish the interaction of MC1 with negatively charged phospholipid head groups. The structure of CtMC1 revealed a large positive patch on the CtMon1 ‘localization layer’ that likely represents the interaction site (Fig S2A). We generated two charge inversion mutants (CIMs) of CtMon1, which had three basic residues exchanged to glutamate in either LD2 (CtMon1^LD2CIM^: R402E, R404E, R551E) or LD3 (CtMon1^LD3CIM^: R636E, R645E, R652E) (Fig 3, S2A). In sedimentation assays, both mutants showed loss of specific binding to PIP-containing liposomes (Fig 4A,B), confirming the model that the basic surface of Mon1 is responsible for the interaction with charged membranes.

**Figure 3:**
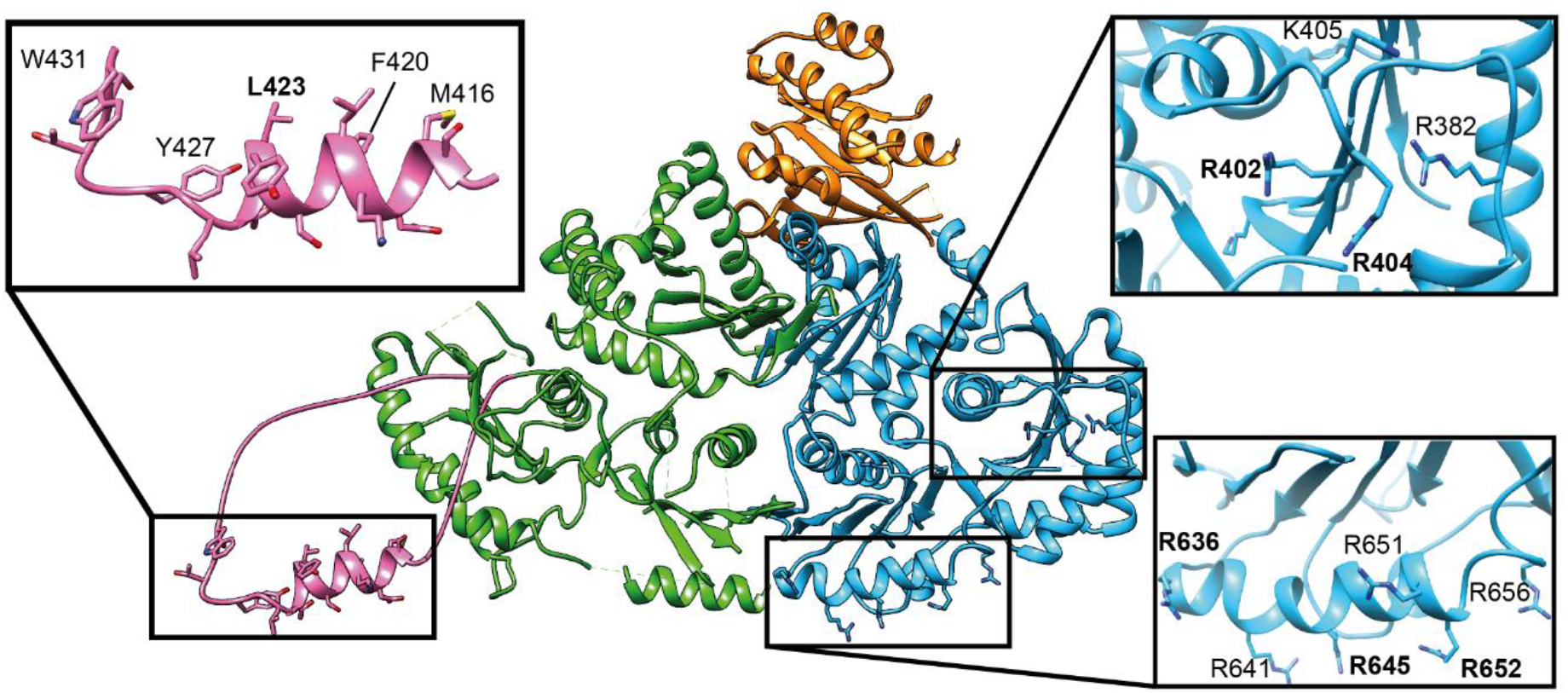
Structural model of the Ct Mon1 (blue)-Ccz1(green) complex bound to Ypt7 (orange). Inserts show the residues contributing to positive charged patches on LD2 and LD3 of Mon1 and the putative amphipathic helix (pink) in the α2-β3 loop of CtCcz1 LD2 as modelled by AlphaFold2 (28), respectively. Mutated residues in CtMon1 and CtCcz1 variants are labeled bold.

**Figure 4:**
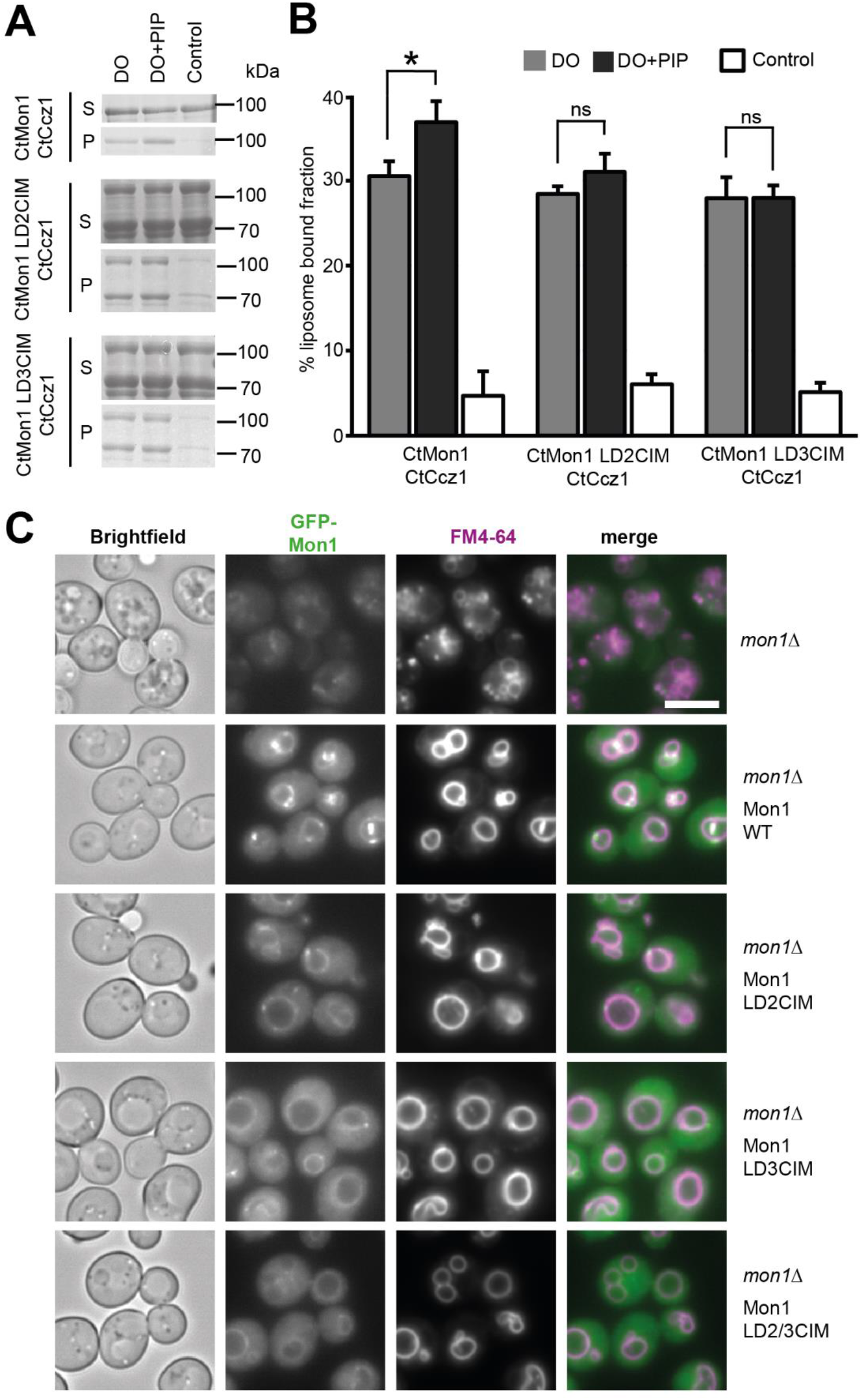
Interaction of the MC1 complex with charged lipids. (**A**) Sedimentation assays of full-length CtMC1 with CtMon1^WT^ or CtMon1^CIM^ variants. Liposomes were generated from a DO lipid mix with or without PIP. (**B**) Quantification of B (n=3). Data are presented as mean ±SD and the significance was calculated using Student’s t-test (* p<0.05, n.s. not significant). (**C**) Fluorescence microscopy images of *mon1*Δ yeast complemented with GFP-ScMon1^WT^ or different GFP-ScMon1^CIM^ variants. Vacuoles are stained with FM4-64. Scale bars: 5 μm.

In analogy, we created mutants of ScMon1 that had charge inversions in LD2 (ScMon1^LD2CIM^: R374E, R376E), LD3 (ScMon1^LD3CIM^: K620E, K624E, K631E) and a combination of both (ScMon1^LD2/3CIM^: R374E, R376E, K620E, R624E, K631E). We introduced these variants as GFP fusion proteins into a *mon1*Δ yeast strain. In the absence of Mon1, yeast cells have fragmented vacuoles (Fig 4C). This phenotype was rescued by the re-introduction of ScMon1^WT^, which localized to perivacuolar puncta that likely represent late endosomes and the vacuolar membrane. Expression of the charge inversion mutants was able to restore vacuolar morphology. However, localization of ScMon1 was impaired (Fig 4C). Perivacuolar puncta were strongly reduced for ScMon1^LD2CIM^ and ScMon1^LD3CIM^ and completely absent for ScMon1^LD2/3CIM^, while some vacuolar membrane staining was still observed. Thus, the basic patch on Mon1 is required for proper localization of the protein to endosomal structures.

To investigate the functionality of ScMon1^CIM^ mutants in autophagy, we introduced the respective plasmids into a *mon1*Δ yeast strain expressing mCherry-Atg8. During starvation, mCherry-Atg8 was properly delivered to the vacuole (Fig S2B), showing that autophagy was not impaired in these cells.

### Identification of a conserved amphipathic helix in Ccz1

We next asked for the possible molecular basis of the interaction of Ccz1 with lipid packing defects. Interestingly, the truncated CtMC1Δ complex we had used for cryo-EM studies was unable to recognize packing defects (Fig S3A,B). Compared to the complexes we used in the mapping experiments (Fig 1), a predicted loop in the LD2 of CtCcz1 (between helix α2 and strand β3, residues 360-460) was removed. Secondary structure analysis and structure predictions of CtCcz1 suggest that this loop contains an α helix with amphipathic character (residues 416-427) (27–29) (Fig 3). Ccz1 homologs from other species also contain a large insertion with a predicted amphipathic helix between α-helix 2 and β-strand 3 of LD2, indicating a functional significance of this motif (Fig 5A). To test if the putative amphipathic helix of Ccz1 can serve as a lipid packing sensor, we generated several CtCcz1 mutants. While CtCcz1^WT^ robustly bound to DO liposomes with packing defects, deletion constructs without the α2-β3 loop (CtCcz1^ΔLoop^, lacking residues 361-460) and without the putative amphipathic helix (CtCcz1^ΔAH^, lacking residues 408-440) lost this ability (Fig 5B,C). Also, introduction of a negative charge in the hydrophobic face of the helix (CtCcz1^L423E^) led to loss of interaction with DO liposomes (Fig 5B,C), as to be expected for a lipid packing sensor function of this helix. A similar effect was also observed with additional point mutants F420E and W431E (Fig S3C).

**Figure 5:**
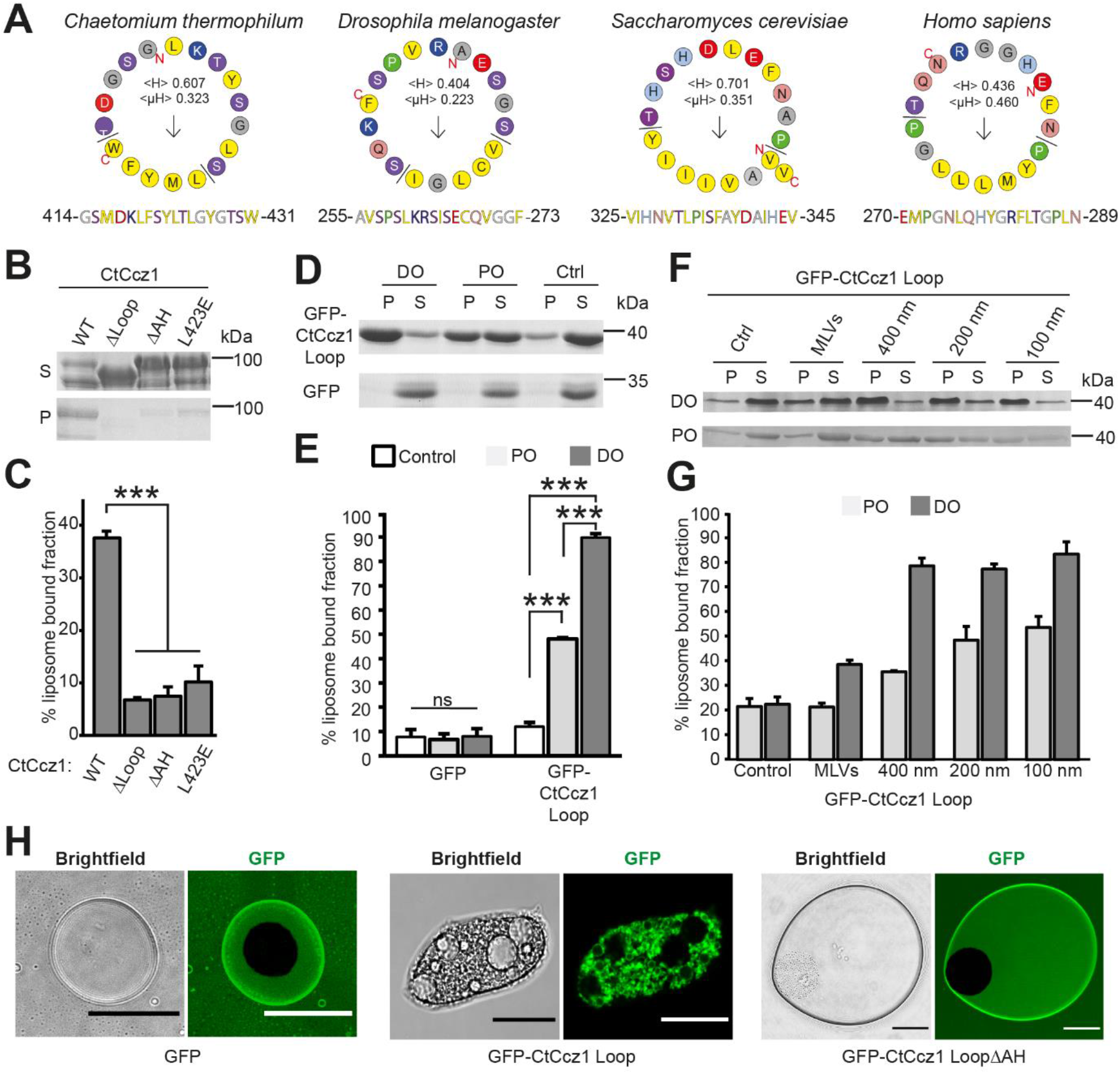
Interaction of the MC1 amphipathic helix with lipids. (**A**) Helical wheel projection of the putative amphipathic helix from different model organisms (29). (**B**) Sedimentation assays of full-length CtCcz1^WT^ or CtCcz1 variants. Liposomes were generated from a DO lipid mix (**C**) Quantification of B (n=3). (**D**) Sedimentation assays of GFP-CtCcz1^Loop^ and GFP with liposomes generated from a PO and DO lipid mix. (**E**) Quantification of D (n=3). (**F**) Sedimentation assays of GFP-CtCcz1^Loop^ with multilamellar vesicles (MLVs) or liposomes with a defined diameter generated from a PO and DO lipid mix. (**G**) Quantification of F (n=4). (**H**) Oil droplets were incubated with GFP, GFP-CtCcz1^Loop^ or GFP-CtCcz1^LoopΔAH^ variants and visualized by confocal fluorescence microcopy. Scale bar: 20 μm. Quantification data are presented as mean ±SD and the significance was calculated using Student’s t-test (*** p<0.001, n.s. not significant).

A fusion of the α2-β3 loop to GFP (GFP-CtCcz1^Loop^, residues 360-460) strongly interacted with DO liposomes, and to lesser extend with PO liposomes that have fewer packing defects (Fig 5D,E). GFP alone did not interact with liposomes. Thus, the α2-β3 loop is not only required, but also sufficient for binding to lipid packing defects. For both PO and DO lipid mixes, curvature-induced lipid packing defects also increased liposome binding of GFP-CtCcz1^Loop^ (Fig 5F,G). Finally, we also tested the effect of lipid packing defects created by different phospholipid head group geometries by including 20% phosphatidylethanolamine (PE, induces negative curvature) or 20% phosphatidylinositol (PI, induces positive curvature) in phosphatidylcholine (PC, no curvature induction) liposomes. In this case, binding of GFP-CtCcz1^Loop^ to liposomes was not changed (Fig S3D,E).

To confirm the amphipathic helix properties of CtCcz1^Loop^, we incubated the GFP fusion protein and GFP alone with an oil suspension (Fig 5H). GFP-CtCcz1^Loop^, but not GFP, acted as an emulsifier of oil droplets, causing the consumption of oil drops into small droplets. The resulting droplets were covered with GFP-CtCcz1^Loop^, showing that the protein has surfactant properties and interacts at the oil-water interface with the hydrophobic surface of the droplets. This behavior is typically observed for amphipathic helices (30). Deletion of the putative amphipathic helix in the Ccz1 loop abolished its ability to emulsify oil droplets. Taken together, these data demonstrate that an amphipathic helix in the α2-β3 loop of Ccz1 LD2 acts as a lipid packing sensor.

### Ccz1 amphipathic helix is required for MC1 function in autophagy

The sedimentation assays with simplified lipid mixes clearly demonstrated the ability of Ccz1 to bind to lipid packing defects, but we wondered if this mechanism would be relevant with more complex lipid mixes that reflect the composition of the target organelles of MC1. Based on lipidomics data of the phospholipid distribution in endosomes and autophagosomes (31–33), we created liposomes that mimic the endosomal and autophagosomal phospholipid composition. These mixtures recapitulate the reported distribution of acyl chains with different degrees of saturation as well as different head group (Fig S3F, Table S1). ScMC1 bound to both lipid mixes, suggesting that the phospholipid compositions of endosomes and autophagosomes in principle generate enough lipid packing defects to support MC1 membrane recruitment (Fig 6A,B). Interestingly, ScMC1 interacted significantly stronger with the autophagosomal lipid mix, which contains acyl chains with more double bonds and thus has more packing defects. Although these differences are subtle, they apparently suffice to modulate membrane binding. When PIPs were included in the liposomes, ScMC1 recruitment could be further enhanced (Fig 6A,B). The difference between the endosomal and autophagosomal lipid mix, however, was diminished under these conditions.

**Figure 6:**
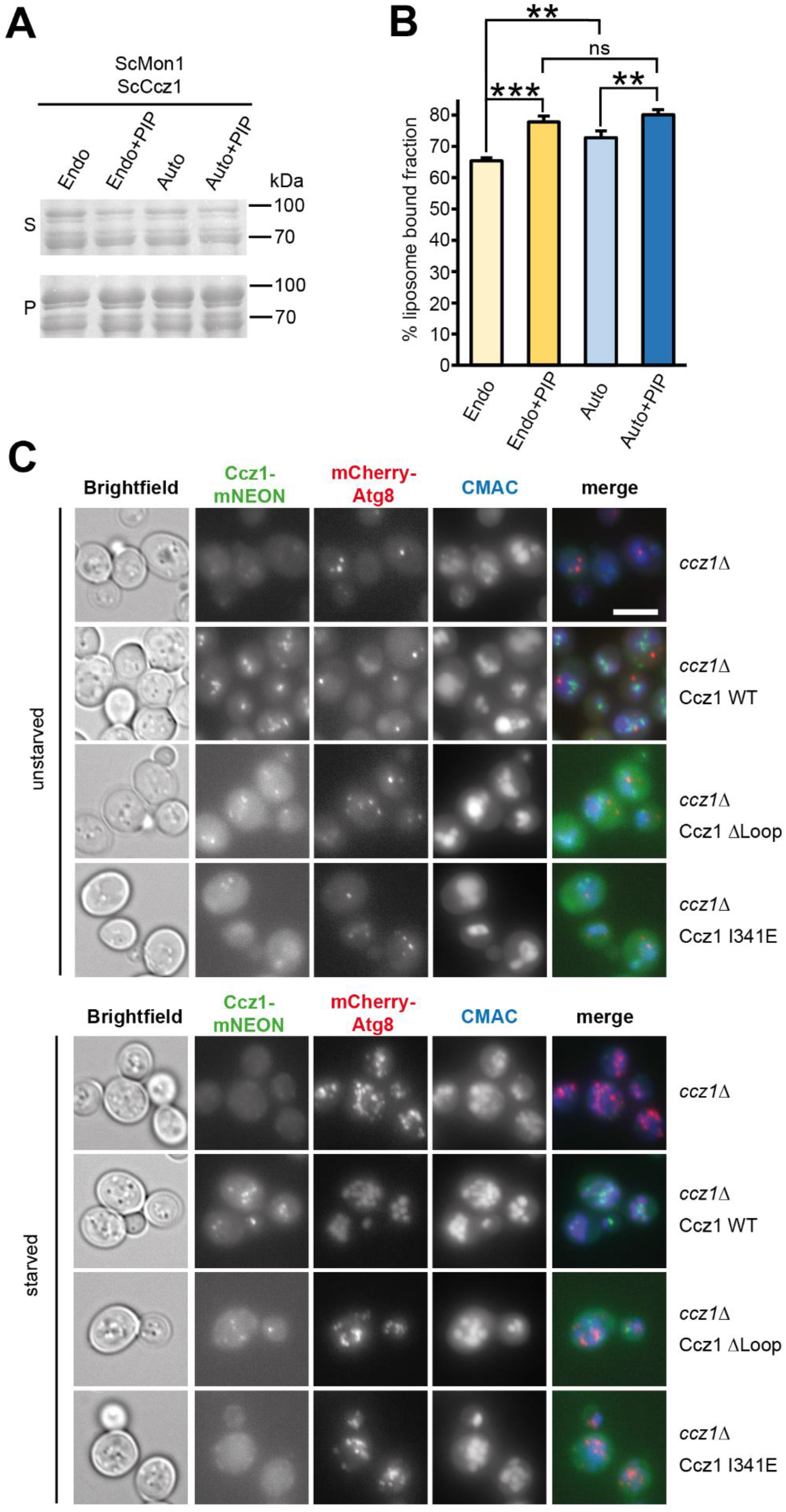
The MC1 complex amphipathic helix is required for autophagy. (**A**) Sedimentation assays of ScMC1 with liposomes generated from an endosomal (endo) or autophagosomal (auto) lipid mix with or without PIP. (**B**) Quantification of B (n=4). Data are presented as mean ±SD and the significance was calculated using Student’s t-test (** p<0.01, *** p<0.001, n.s. not significant). (**C**) Immunofluorescence images of unstarved and starved *ccz1*Δ yeast expressing mCherry-Atg8 and complemented with GFP-ScCcz1^WT^ or different GFP-ScCcz1 variants. Vacuoles are stained with CMAC. Scale bar: 5 μm.

For functional testing, we generated in analogy to CtCcz1 a loop deletion mutant ScCcz1^ΔLoop^ (lacking residues 270-403) and a variant ScCcz1^I341E^, which has a charged residue introduced in the hydrophobic face of the amphipathic helix. Loss of Ccz1 in yeast caused vacuolar fragmentation, which was rescued by reintroduction of ScCcz1^WT^. The same effect was observed with ScCcz1^ΔLoop^ and ScCcz1^I341E^, which localized like ScCcz1^WT^ to punctate structures (Fig 6C). Thus, the amphipathic helix is not required for endosomal recruitment of ScMC1 or endosomal maturation. However, when we investigated the functionality of ScCcz1 mutants in starved cells, an autophagy defect was observed. Like in *ccz1*Δ cells, Atg8 did not reach the vacuole after complementation with ScCcz1^ΔLoop^ and ScCcz1^I341E^, whereas ScCcz1^WT^ was able to restore autophagic flux (Fig 6C). The ScCcz1^ΔLoop^ and ScCcz1^I341E^ strains also showed a defect in Ape1 processing, an assay monitoring the uptake of oligomeric Ape1 peptidase via the autophagy-related cytosol-to-vacuole targeting (CVT) pathway (34) (Fig S3G). The data therefore show that the amphipathic helix is necessary for the function of ScMC1 in autophagy.

## Discussion

The identification of an amphipathic helix of the MC1 complex that mediates binding to membranes by recognizing lipid packing defects as a novel targeting mechanism by which the complex is recruited to its proper localization in the cell. This activity is conserved and could be observed for both CtMC1 and ScMC1. A putative amphipathic helix motif is also found in other Ccz1 homologs at the same position in the α2-β3 loop of LD2. Thus, the lipid packing sensing function may also play a role in other species.

Lipid packing defects act synergistically with negative charges of phospholipid head groups and recruiter proteins in recruitment of MC1 to model membranes. While each interaction individually is relatively weak, two targeting cues together strongly promote membrane binding. A close to stoichiometric recruitment, however, was only achieved if all three cues were present. This behavior is consistent with independent binding events where affinities are multiplicative rather than additive. Alternatively, the observed effects could potentially also arise, in parts, from cooperativity between binding events.

The observed synergistic binding could be the basis for an ultrasensitive behavior in complex recruitment that encodes spatiotemporal regulation (23, 24). In particular, the levels of PIPs and recruiter proteins accumulate during the maturation of both endosomes and autophagosomes (25, 26). It is thus tempting to speculate that coincidence detection employing multiple targeting mechanisms with individually weak affinities allows for tightly controlled, yet rapid recruitment and activation of the MC1 GEF and the downstream pathways when thresholds are reached.

The simplified liposome mixes we used for our *in vitro* studies do not fully recapitulate the complex properties of biological membranes with membrane proteins, sterols or sphingolipids. Thus, the different targeting cues may not all be required (to the same extend) for all functions of MC1. In fact, our functional analyses indicate that binding to charged lipids is required for endosomal localization of ScMC1 and interaction with packing defects for its function in autophagy, but not vice versa. This is consistent with the concept that protein targeting to endosomal organelles is dominated by charges, while targeting to secretory compartments (autophagosomes receive their bulk lipids from the ER (35)) is dominated by packing defect (36). By exploiting distinct membrane properties for differential recruitment, MC1 targeting is different from the mechanism that drives localization of the Rab1 GEF TRAPPIII to the Golgi and autophagosomes. TRAPPIII also requires binding to packing defects and charges for proper function, but both cues are needed at both sites action (37). It is interesting to note that the endosomal Rab5 GTPases bind Mon1 (15, 18) and the autophagosome marker Atg8 binds Ccz1 (11). This suggests a model were Mon1 and Ccz1 together are required for catalytic activity of the complex (22), but each subunit is responsible to mediate a distinct localization through the recognition of unique protein and lipid cues (Fig 7).

**Figure 7:**
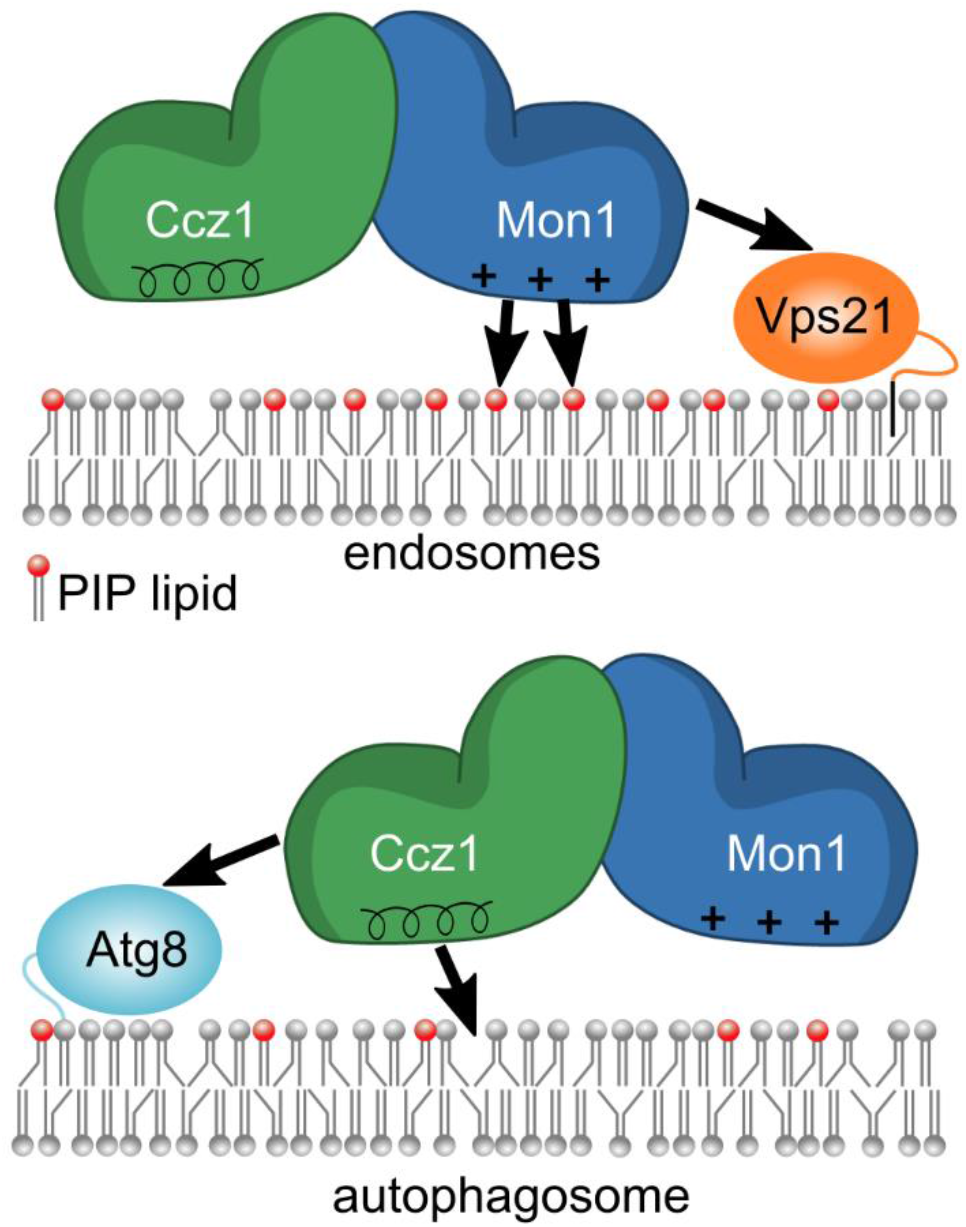
Model for the differential targeting of MC1 to endosomes and autophagosomes. Mon1 interacts with Rab5 family GTPases (Vps21) and PIPs to recruit the complex to endosomes. For binding to autophagosomes, Ccz1 instead interacts with Atg8 and lipid packing defects.

Thus, the complex has adapted to the different requirements for targeting in distinct pathways: MC1 functions on endosomes, which have high PI3P levels that are produced during endosomal maturation. This membrane property is exploited by Mon1 through charge interactions. In contrast, MC1 acts early during autophagosome formation when PIP levels are low and the requirement for recognition of an alternative lipid cue arises. Similar to other autophagosomal proteins that employ amphipathic helices for recruitment (38), Ccz1 utilizes a membrane packing/curvature sensing motif for association with the phagophore. Our findings support the concept of a mechanistic role for the high degree of unsaturation in autophagosomal phospholipid acyl chains observed before (32).

The targeting of MC1 can likely also be fine-tuned by post-translational modifications, e.g. phosphorylation that was reported for Mon1 (15), which may promote or inhibit targeting. Importantly, distinct protein and lipid binding mechanisms to endosomes and autophagosomes allow differential and independent regulation of both MC1 functions. We expect that this concept may also be realized by other regulatory trafficking complexes that serve multiple functions in the cell.

## Experimental procedures

### Protein expression and purification

CtMon1 constructs were cloned into pCDF6P (N-terminal GST-tag, PreScission protease cleavage site) and CtCcz1 constructs into pET28HS (N-terminal 6xHis-SUMO tag) (Table S2). *E. coli* BL21 (DE3) cells were co-transformed with different construct combinations, or CtCcz1 alone, and expression was induced after cold shock with 0.25 mM isopropyl-β-D-thiogalactoside for 16 h shaking at 16°C. Cells were lyzed in buffer A (50 mM NaH_2_PO_4_, 500 mM NaCl, 1 mM MgCl_2_, 5% (v/v) glycerol, pH 7.3) with protease inhibitor mix HP (Serva). Cleared supernatants were incubated with glutathione agarose, beads were washed and proteins were eluted by sequential proteolytic cleavage of the tags with SUMO protease (2h) and PreScission protease (overnight), or by addition of with 20 mM glutathione and 12 mM DTT. Eluates were further purified by size exclusion chromatography (SEC650, Bio-Rad) pre-equilibrated with buffer B (25 mM HEPES, 250mM NaCl, 1mM MgCl_2_, 0.5 mM TCEP, pH 7.3). CtCcz1 single expressions were purified via Ni-NTA affinity chromatography, elution with 250 mM imidazole and size exclusion chromatography as described above.

ScMC1 was expressed under the *GAL1* promotor in yeast (CUY2470: BY4732; *CCZ1::TRP1-GAL1pr MON1::HIS3MX6-GAL1pr CCZ1::TAP-URA3*) (6). Culture were grown in YP+galactose media at 30°C to OD600 of 5 and harvested by centrifugation. Cells were lyzed in buffer Y (50 mM HEPES-NaOH, pH 7.4, 150 mM NaCl, 1.5 mM MgCl_2_, FY protease inhibitor mix (Serva), 0.5 mM PMSF, 1 mM DTT). Cleared lysates were incubated with IgG Sepharose (GE Healthcare), washed with buffer Y and eluted by overnight TEV cleavage.

GST-TEV-Vps21 and GST-TEV-Ypt10 were purified with glutathione affinity chromatography, on-column TEV-protease cleavage overnight and dialyzed against assay buffer (50 mM HEPES, NaOH pH 7.4, 150 mM NaCl, 1 mM MgCl_2_) (15). His-TEV-Atg8 was purified via Ni-NTA affinity chromatography, followed by TEV cleavage in solution, cation exchange chromatography and gel filtration (running buffer 50 mM HEPES pH 7.5, 150 mM NaCl, 1 mM DTT) (39).

### Liposome preparation

Lipids (Avanti Polar Lipids, except PIPs purchased from Echelon Biosciences) were mixed in chloroform and dried in a speedvac. The compositions of the different lipid mixes used are listed in Table S1. The lipid film was dissolved in 1 ml of buffer L (25 mM HEPES, 250 mM NaCl, 1 mM MgCl_2_, 5% sucrose, pH 7.3) to a final lipid concentration of 2 mM. Multilamellar lipid vesicles were generated by five cycles of freezing in liquid nitrogen and thawing at 56°C and stored at −80 °C. Prior to use, MLVs were extruded 21 times through a polycarbonate membrane to generate liposomes of 400 nm diameter unless stated otherwise.

### Prenylation of Vps21 and Ypt10

Prior to prenylation, GTPases (40 μM) were loaded with GDP (80 μM) by incubation in the presence of 20 mM EDTA for 30 minutes at 30°C, followed by the addition of 25 mM MgCl_2_. The prenylation was performed according to a protocol modified from (40). Each reaction contained 3 μM GTPase (preloaded with GDP), 3 μM Mrs6 (Rab escort protein), 1 μM Bet2/Bet4 (Rab-GGtase), 9 μM geranylgeranylpyrophosphate in buffer P (20 mM HEPES, 150 mM NaCl, 1 mM DTT, pH 7.3) was incubated for 1h at 37°C and samples were stored at −80°C. To bind prenylated GTPases onto liposomes, prenylated GTPases were incubated with liposomes for 20 min at room temperature in the presence of 1 mM GTP.

### Lipidation of Atg8

MLVs were extruded to generate 400 nm liposomes. Lipidation of Atg8 was preformed essentially as described before (39). 1 mM liposomes were mixed with 1 μM Atg7, 1.25 μM Atg3, 2.5 μM Atg8, 0.5 μM Atg5-12, 0.5 μM Atg16, 2 mM ATP, 1 mM MgCl_2_ and 1 mM DTT (all final concentrations) and incubated for 30 min at 30°C. Samples were flash-frozen and stored at −80°C.

### Liposome sedimentation assays

Proteins and liposomes were mixed in buffer B to final concentrations of 1 μM and 0.5 mM, respectively, in a final volume of 200 μl. Reactions were incubated for 20 min at room temperature and liposomes were pelleted at 20,000 x *g* for 20 min at 4°C. The soluble supernatant fraction was precipitated with acetone on ice and supernatant and pellet fractions were analyzed by SDS-PAGE and Coomassie staining. For quantifications, gel band intensities were measured with ImageJ to calculate the liposome bound fraction of protein. In samples with Mrs6, which migrates at the same molecular weight as Mon1 in SDS-PAGE, only Ccz1 bands were quantified. A two-tailed heteroscedastic t-test was used for significance analyses.

### Oil emulsion assay

Purified GFP, GFP-CtCcz1^Loop^ GFP-CtCcz1^LoopΔAH^ (500 μl of at a concentration of 2 mg ml^−1^) were mixed with 20 μl olive oil (REWE Beste Wahl, extra virgin) and vortexed for 2 min. After incubation at room temperature for 20 min, phase separation was observed with GFP and GFP-CtCcz1^LoopΔAH^. For GFP-CtCcz1^Loop^ formation of an emulsion phase was observed. A 10μl sample of the emulsion or separation phase, respectively, was transferred to a glass slide and covered with a cover slip. Samples were visualized using a DMI-8-CS l microscope (Leica) with HC PL APO UVIS CS2 63×1.20 water or HC PC FLUOTAR 10×0.30 dry objective lens and the PMT trans detector. Images were acquired using the Leica software and prepared with ImageJ.

### Yeast strains

The yeast strains used in this study carry a deletion of either Mon1 or Ccz1 and express mCherry-Atg8 under the control of the ADH promotor (CUY10469 mCherry-Atg8 *ccz1*Δ (MATalpha *leu2-3,112 ura3-52 his3*-Δ*200 trp*-Δ*901 lys2-801 suc2*-Δ*9 GAL ATG8::ADHpr-mCherry-natNT2 ccz1*Δ*::hphNT1*) and CUY13522 mCherry Atg8 *mon1*Δ (MATalpha *leu2-3,112 ura3-52 his3*-Δ*200 trp*-Δ*901 lys2-801 suc2*-Δ*9 GAL MON1::hphNT1 ATG8::ADHpr-mCherry-natNT2*)). ScMon1 variants were cloned into a pRS406 plasmid with an N-terminal GFP-tag under the control of the NOP1 promotor. ScCcz1 and variants were cloned into a pRS406 plasmid with a C-terminal mNEON-tag under the control of the CCZ1 promotor. The plasmids (Table S2) were linearized and transformed into the respective *mon1*Δ or *ccz1*Δ strains by integration into the *URA*3 locus, respectively.

### Microscopy of starved cells

Cells were grown in yeast extract peptone medium containing glucose (YPD) overnight, diluted to OD_600_ of 0.2 in the morning and grown for 2.5h. Cells were washed either in minimal synthetic medium lacking nitrogen (SD-N) or in YPD and then grown for another 2h in the respective medium before preparation for microscopy. Cells were collected by centrifugation (5,000 *g*, 3 min, 20 °C) and washed in synthetic media once. For staining of the vacuole by CMAC, cells were incubated in synthetic media with or without nitrogen source containing 0.1 μM CMAC for 15 min at 30 °C, washed twice in fresh media and incubated another 15 min in media without dye. Images were acquired directly afterwards at room temperature using a Delta Vision Elite (GE Heathcare) equipped with an inverted microscope (model IX-71; Olympus), an UAPON X 100 (1.49 numerical aperture (NA)) oil immersion, an InsightSSI light source (Applied Precision) and an sCMOS camera (PCO). Data were processed using ImageJ 2.1.0. Shown pictures are maximum intensity projections of medial planes of yeast cells.

### Microscopy of Mon1^CIM^ mutants

Cells were grown in yeast extract peptone medium containing glucose (YPD) overnight, diluted to OD_600_ of 0.25 in the morning and grown until an OD_600_ of around 1. Cells were collected by centrifugation (5,000 *g*, 3 min, 20 °C) and washed in synthetic media once. For staining of the vacuole by FM4-64, cells were incubated in synthetic media containing 30 μM FM4-64 for 30 min at 30 °C, washed twice in fresh media, and incubated another 45 min in media without dye. Images were acquired directly afterwards using a Zeiss Axioscope 5 FL (Zeiss) equipped with Plan-Apochromat 100x (1.4 numerical aperture (NA)) oil immersion objective and an Axiocam 702 mono camera. Data were obtained using the ZEN 3.1 pro software and processed using ImageJ 2.1.0. Shown pictures are single medial planes of yeast cells.

### Ape1 processing assay

Cells were grown overnight in YPD medium, diluted in the morning to an OD_600_ of 0.3 and grown until an OD_600_ of around 1. They were washed either in SD-N or in YPD media and then incubated for another 90min. Cells corresponding to 2 OD600 were harvested, resuspended in 500 μl ice-cold H_2_O and lysed by immediate addition of 75 μl 1,85 M NaOH containing 1M beta-mercaptoethanol and 1 mM PMSF. After incubation for 10 min at 4°C a final concentration of 13% (v/v) trichloroacetic acid was added and incubated again on ice for 15 min. Samples were centrifuged for 15 min at 20,000 *g* and pellet was washed in 1 ml ice-cold acetone followed by centrifugation for 15 min at 20,000 *g*. Pellet was dried and applied to SDS-PAGE and Western-blot for further analysis by a polyclonal antibody directed against Ape1.

## Data availability

All data supporting the findings of this study are available from the corresponding author upon request.

## Supporting Information

This article contains supporting information.

## Acknowledgments

We are grateful to Carmen Gelze and Jesse Tönjes for excellent technical assistance and to Abdou Rachid Thiam and Katharina Höffgen for helpful discussions. We thank Sascha Martens (University of Vienna, Austria) for providing us with the lipidation machinery for Atg8.

## Author contributions

Investigation and methodology: E. Herrmann, L. Langemeyer. Conceptualization and writing – first draft: D. Kümmel. Writing – review & editing: E. Herrmann, L. Langemeyer, C. Ungermann. Funding acquisition: C. Ungermann, D. Kümmel

## Supporting information

**Figure S1:**
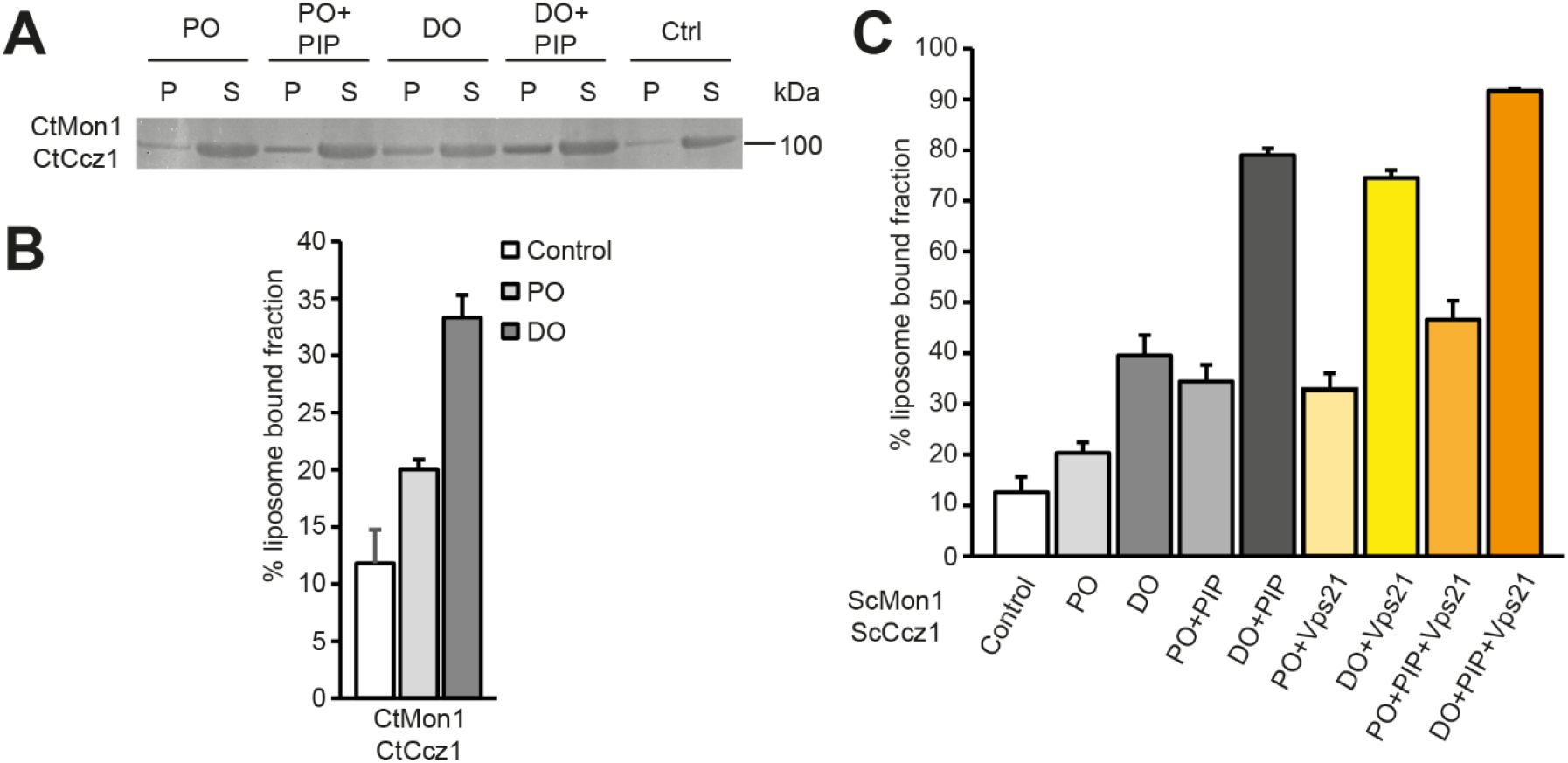
Lipid binding properties of MC1. (**A**) Sedimentation assays of full-length CtMC1 with liposomes form a PO (palmitoyl-oleoyl) and DO (di-oleoyl) lipid mix (**B**) Quantification of A (n=4). (**C**) Quantification of binding of ScMC1 to liposomes without and with Vps21, high packing defects (DO), charged lipids (PIP), and different combinations (n=3-8). The graph summarizes data presented in Figures 2.

**Figure S2:**
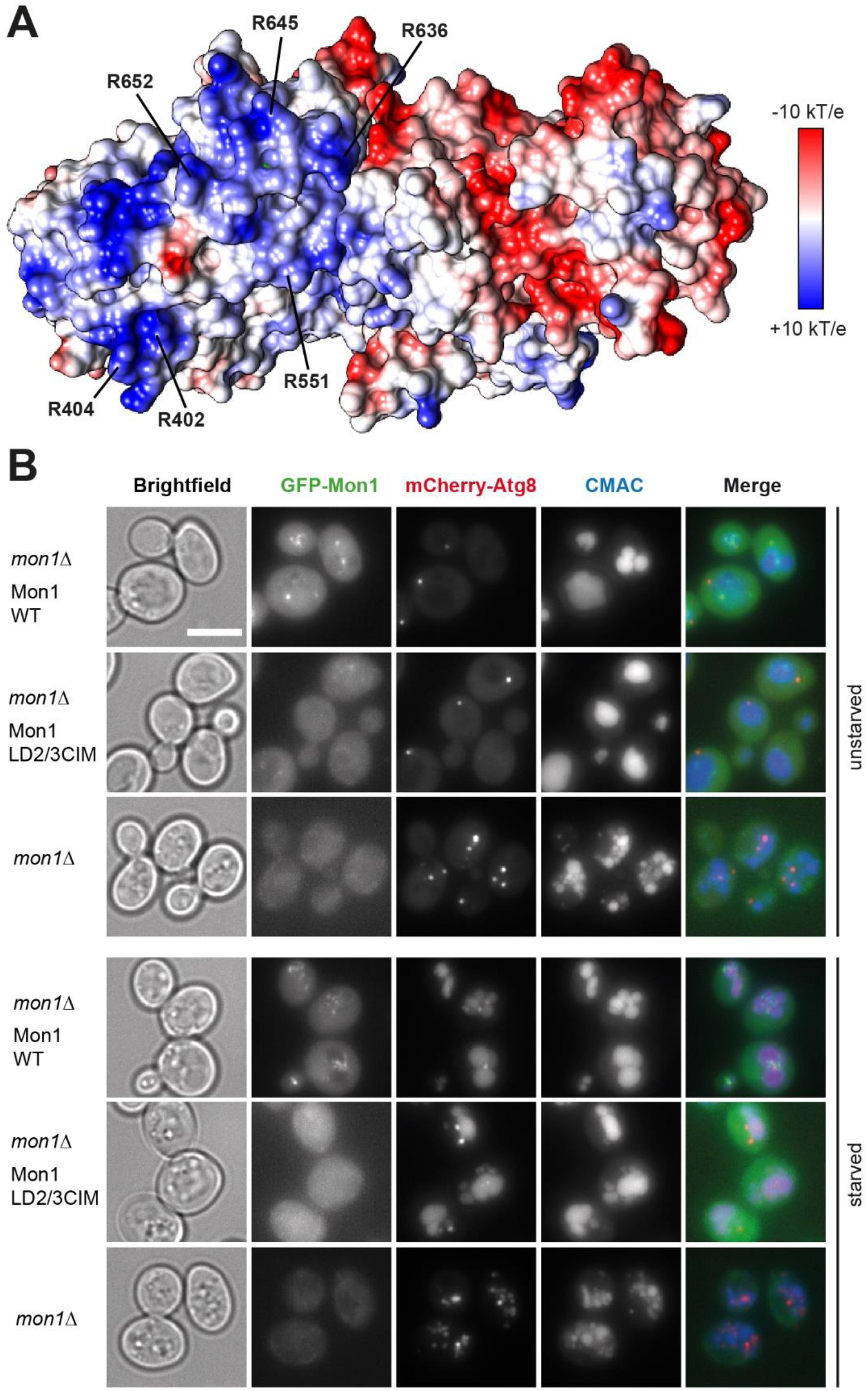
Role of a charged patch on Mon1 in MC1 complex function. (**A**) Surface potential representation of the putative membrane binding interface of the CtMC1 complex. The locations of key basic residues of Mon1 involved in PIP binding are indicated. (**B**) Fluorescence microscopy images of *mon1*Δ yeast expressing mCherry-Atg8 and complemented with GFP-ScMon1^WT^ or different GFP-ScMon1^CIM^ variants. Vacuoles are stained with CMAC.

**Figure S3:**
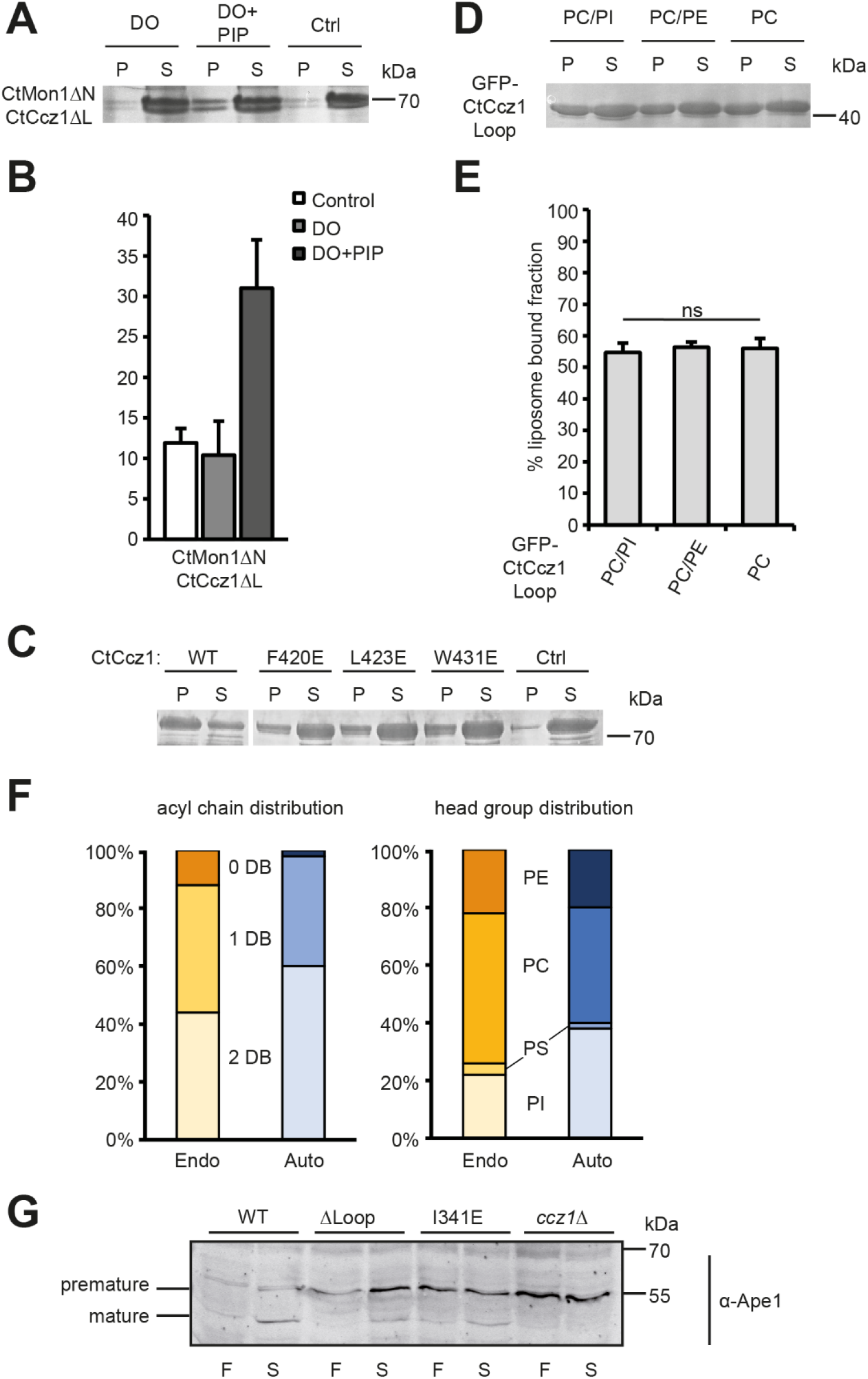
Characterization of the Ccz1 amphipathic helix. (**A**) Sedimentation assays of CtMC1Δ (CtMon1 141-665/CtCcz1 1-360,461-796) with liposomes form DO (di-oleoyl) lipid mix with or without PIP (**B**) Quantification of A (n=4). (**C**) Reduced binding of CtCcz1 point mutants in the amphipathic helix to DO liposomes. (**D**) Binding of GFP-CtCcz1^Loop^ to di-oleoyl phosphatidylcholine (PC) with or without 20% di-oleoyl phosphatidylinositol (PI) or di-oleoyl phosphatidylethanolamine (PE). (**E**) Quantification of D (n=3). Quantification data are presented as mean ±SD. (**F**) Phospholipid composition of yeast endosomes (endo) and autophagosomes (auto). DB: double bond; PI: phosphatidylinositol; PS: phosphatidylserine; PC: phosphatidylcholine; PE: phosphatidylethanolamine. (**G**) Ape1 processing assay of fed (F) and starved (S) *ccz1*Δ yeast complemented with GFP-ScCcz1^WT^ or different GFP-ScCcz1 variants.

**Table S1:** Composition of lipid mixes used in this study

| **Lipid mix** | **Composition** |
| --- | --- |
| PO | 82 mol% 1-palmitoyl-2-oleoyl phosphatidylcholine<br>18 mol% 1-palmitoyl-2-oleoyl phosphatidylethanolamine |
| PO+PIP | 79 mol% 1-palmitoyl-2-oleoyl phosphatidylcholine<br>18 mol% 1-palmitoyl-2-oleoyl phosphatidylethanolamine<br>2 mol% dipalmitoylphosphatidylinositol-3-phosphate<br>1 mol% dipalmitoylphosphatidylinositol-3,5-bisphosphate |
| DO | 82 mol% dioleoyl phosphatidylcholine<br>18 mol% dioleoyl phosphatidylethanolamine |
| DO+PIP | 79 mol% dioleoyl phosphatidylcholine<br>18 mol% dioleoyl phosphatidylethanolamine<br>2 mol% dipalmitoylphosphatidylinositol-3-phosphate<br>1 mol% dipalmitoylphosphatidylinositol-3,5-bisphosphate |
| PC | 100 mol% 1-palmitoyl-2-oleoyl phosphatidylcholine |
| PC/PI | 80 mol% 1-palmitoyl-2-oleoyl phosphatidylcholine<br>20 mol% 1-palmitoyl-2-oleoyl phosphatidylinositol |
| PC/PE | 80 mol% 1-palmitoyl-2-oleoyl phosphatidylcholine<br>20 mol% 1-palmitoyl-2-oleoyl phosphatidylethanolamine |
| Endo | 40 mol% dioleoyl phosphatidylcholine<br>12 mol% dipalmitoyl phosphatidylcholine<br>22 mol% 1-palmitoyl-2-oleoyl phosphatidylethanolamine<br>22 mol% 1-palmitoyl-2-oleoyl phosphatidylinositol<br>4 mol% dioleoyl phosphatidylserine |
| Auto | 38 mol% dioleoyl phosphatidylcholine<br>2 mol% dipalmitoyl phosphatidylcholine<br>20 mol% dioleoyl phosphatidylethanolamine<br>38 mol% 1-palmitoyl-2-oleoyl phosphatidylinositol<br>2 mol% dioleoyl phosphatidylserine |
| Endo+PIP | 40 mol% dioleoyl phosphatidylcholine<br>2 mol% 1-palmitoyl-2-oleoyl phosphatidylcholine<br>10 mol% dipalmitoyl phosphatidylcholine<br>22 mol% 1-palmitoyl-2-oleoyl phosphatidylethanolamine<br>20 mol% 1-palmitoyl-2-oleoyl phosphatidylinositol<br>4 mol% dioleoyl phosphatidylserine<br>2 mol% dipalmitoylphosphatidylinositol-3-phosphate |
| Auto+PIP | 38 mol% dioleoyl phosphatidylcholine<br>2 mol% 1-palmitoyl-2-oleoyl phosphatidylcholine<br>20 mol% dioleoyl phosphatidylethanolamine<br>36 mol% 1-palmitoyl-2-oleoyl phosphatidylinositol<br>2 mol% dioleoyl phosphatidylserine<br>2 mol% dipalmitoylphosphatidylinositol-3-phosphate |

**Table S2:** Plasmids used in this study

| Plasmid | Construct |
| --- | --- |
| DK65 | pCDF6P-CtMon1<sup>FL</sup><br>(GST-PreScission site CtMon1 1-665) |
| DK44 | pCDF6P-CtMon1<sup>ΔN</sup><br>(GST-PreScission site CtMon1 141-665) |
| DK123 | pCDF6P-CtMon1<sup>LD1</sup><br>(GST-PreScission site CtMon1 141-355) |
| DK1534 | pCDF6P-CtMon1<sup>LD2CIM</sup><br>(GST-PreScission site CtMon1 141-665; R402E, R404E, R551E) |
| DK1535 | pCDF6P-CtMon1<sup>LD3CIM</sup><br>(GST-PreScission site CtMon1 141-665; R636E, R645E, R652E) |
| CU3481 | pRS406-GFP-ScMon1<sup>WT</sup><br>(NOP1pr GFP-ScMon1 1-644) |
| DK1585 | pRS406-GFP-ScMon1<sup>LD2CIM</sup><br>(NOP1pr GFP-ScMon1 1-644; R374E, R376E) |
| DK1514 | pRS406-GFP-ScMon1<sup>LD3CIM</sup><br>(NOP1pr GFP-ScMon1 1-644; K620E, K624E, K631E) |
| DK1585 | pRS406-GFP-ScMon1<sup>LD2/3CIM</sup><br>(NOP1pr GFP-ScMon1 1-644; R374E, R376E, K620E, R624E, K631E) |
| DK46 | pET28HS-CtCcz1<sup>FL</sup><br>(6xHis-SUMO CtCcz1 1-796) |
| DK48 | pET28HS-CtCcz1<sup>LD1</sup><br>(6xHis-SUMO CtCcz1 1-249) |
| DK857 | pET28HS CtCcz1<sup>ΔLoop</sup><br>(6xHis-SUMO CtCcz1 1-360,461-796) |
| DK1052 | pET28HS CtCcz1<sup>ΔAH</sup><br>(6xHis-SUMO CtCcz1 1-407,441-796) |
| DK1149 | pET28HS CtCcz1<sup>L423E</sup><br>(6xHis-SUMO CtCcz1 1-796; L423E) |
| DK134 | pET28GFP<br>(6xHis-Thrombin site-GFP) |
| DK1287 | pET28GFP-CtCcz1<sup>Loop</sup><br>(6xHis-Thrombin site-GFP CtCcz1 360-460) |
| DK1677 | pET28GFP-CtCcz1<sup>LoopΔAH</sup><br>(6xHis-Thrombin site-GFP CtCcz1 360-407,441-460) |
| CU5525 | pRS406-ScCcz1<sup>WT</sup>-mNEON<br>(CCZ1pr-ScCcz1 1-704 mNEON) |
| CU5528 | pRS406-ScCcz1<sup>ΔLoop</sup>-mNEON<br>(CCZ1pr-ScCcz1 1-269,404-704 mNEON) |
| CU5527 | pRS406-ScCcz1<sup>I341E</sup>-mNEON<br>(CCZ1pr-ScCcz1 1-704; I341E mNEON) |
| CU2452 | pET24d-GST-TEV-Vps21 |
| CU4527 | pET24d GST-TEV-Ypt10 |
| CU5326 | pET-DUET-1-HIS-TEV-Atg8 (Gift from Sascha Martens, (39)) |

## References

1. Holthuis, J. C. M., and Ungermann, C. (2013) Cellular microcompartments constitute general suborganellar functional units in cells. Biol. Chem. 394, 151–61

2. Ardito, F., Giuliani, M., Perrone, D., Troiano, G., and Muzio, L. Lo (2017) The crucial role of protein phosphorylation in cell signaling and its use as targeted therapy (Review). Int. J. Mol. Med. 40, 271–280

3. Bos, J. L., Rehmann, H., and Wittinghofer, A. (2007) GEFs and GAPs: Critical Elements in the Control of Small G Proteins. Cell. 129, 865–877

4. Carlton, J. G., and Cullen, P. J. (2005) Coincidence detection in phosphoinositide signaling. Trends Cell Biol. 15, 540–547

5. Di Paolo, G., and De Camilli, P. (2006) Phosphoinositides in cell regulation and membrane dynamics. Nature. 443, 651–657

6. Nordmann, M., Cabrera, M., Perz, A., Bröcker, C., Ostrowicz, C., Engelbrecht-Vandré, S., and Ungermann, C. (2010) The Mon1-Ccz1 complex is the GEF of the late endosomal Rab7 homolog Ypt7. Curr. Biol. 20, 1654–9

7. Gerondopoulos, A., Langemeyer, L., Liang, J.-R., Linford, A., and Barr, F. A. (2012) BLOC-3 mutated in Hermansky-Pudlak syndrome is a Rab32/38 guanine nucleotide exchange factor. Curr. Biol. 22, 2135–9

8. Kucharczyk, R., Kierzek, A. M., Slonimski, P. P., and Rytka, J. (2001) The Ccz1 protein interacts with Ypt7 GTPase during fusion of multiple transport intermediates with the vacuole in S. cerevisiae. J. Cell Sci. 114, 3137–45

9. Kucharczyk, R., Dupre, S., Avaro, S., Haguenauer-Tsapis, R., Slonimski, P. P., and Rytka, J. (2000) The novel protein Ccz1p required for vacuolar assembly in Saccharomyces cerevisiae functions in the same transport pathway as Ypt7p. J. Cell Sci. 113, 4301–4311

10. Poteryaev, D., Datta, S., Ackema, K., Zerial, M., and Spang, A. (2010) Identification of the switch in early-to-late endosome transition. Cell. 141, 497–508

11. Gao, J., Langemeyer, L., Kümmel, D., Reggiori, F., and Ungermann, C. (2018) Molecular mechanism to target the endosomal Mon1-Ccz1 GEF complex to the pre-autophagosomal structure. Elife. 7, e31145

12. van den Boomen, D. J. H., Sienkiewicz, A., Berlin, I., Jongsma, M. L. M., van Elsland, D. M., Luzio, J. P., Neefjes, J. J. C., and Lehner, P. J. (2020) A trimeric Rab7 GEF controls NPC1-dependent lysosomal cholesterol export. Nat. Commun. 11, 5559

13. Hegedüs, K., Takáts, S., Boda, A., Jipa, A., Nagy, P., Varga, K., Kovács, A. L., and Juhász, G. (2016) The Ccz1-Mon1-Rab7 module and Rab5 control distinct steps of autophagy. Mol. Biol. Cell. 27, 3132–3142

14. Yan, B.-R., Li, T., Coyaud, E., Laurent, E. M. N., St-Germain, J., Zhou, Y., Kim, P. K., Raught, B., and Brumell, J. H. (2021) C5orf51 is a component of the MON1-CCZ1 complex and controls RAB7A localization and stability during mitophagy. Autophagy. 10.1080/15548627.2021.1960116

15. Langemeyer, L., Borchers, A.-C., Herrmann, E., Füllbrunn, N., Han, Y., Perz, A., Auffarth, K., Kümmel, D., and Ungermann, C. (2020) A conserved and regulated mechanism drives endosomal Rab transition. Elife. 9, e56090

16. Vaites, L. P., Paulo, J. A., Huttlin, E. L., and Harper, J. W. (2017) Systematic Analysis of Human Cells Lacking ATG8 Proteins Uncovers Roles for GABARAPs and the CCZ1/MON1 Regulator C18orf8/RMC1 in Macroautophagic and Selective Autophagic Flux. Mol. Cell. Biol. 38, e00392–17

17. Cabrera, M., Nordmann, M., Perz, A., Schmedt, D., Gerondopoulos, A., Barr, F., Piehler, J., Engelbrecht-Vandré, S., and Ungermann, C. (2014) The Mon1-Ccz1 GEF activates the Rab7 GTPase Ypt7 via a longin-fold-Rab interface and association with PI3P-positive membranes. J. Cell Sci. 127, 1043–51

18. Lawrence, G., Brown, C. C., Flood, B. A., Karunakaran, S., Cabrera, M., Nordmann, M., Ungermann, C., and Fratti, R. A. (2014) Dynamic association of the PI3P-interacting Mon1-Ccz1 GEF with vacuoles is controlled through its phosphorylation by the type 1 casein kinase Yck3. Mol. Biol. Cell. 25, 1608–1619

19. Gerondopoulos, A., Strutt, H., Stevenson, N. L., Sobajima, T., Levine, T. P., Stephens, D. J., Strutt, D., and Barr, F. A. (2019) Planar Cell Polarity Effector Proteins Inturned and Fuzzy Form a Rab23 GEF Complex. Curr. Biol. 29, 3323–3330

20. Sanchez-Pulido, L., and Ponting, C. P. (2019) Hexa-Longin domain scaffolds for inter-Rab signalling. Bioinformatics. 36, 990–993

21. Klink, B. U., Herrmann, E., Antoni, C., Langemeyer, L., Kiontke, S., Gatsogiannis, C., Ungermann, C., Raunser, S., and Kümmel, D. (2022) Structure of the Mon1-Ccz1 complex reveals molecular basis of membrane binding for Rab7 activation. Proc. Natl. Acad. Sci. U. S. A. 119, e2121494119

22. Kiontke, S., Langemeyer, L., Kuhlee, A., Schuback, S., Raunser, S., Ungermann, C., and Kümmel, D. (2017) Architecture and mechanism of the late endosomal Rab7-like Ypt7 guanine nucleotide exchange factor complex Mon1-Ccz1. Nat. Commun. 8, 14034

23. Barr, F. A. (2013) Review series: Rab GTPases and membrane identity: causal or inconsequential? J Cell Biol. 202, 191–199

24. Bezeljak, U., Loya, H., Kaczmarek, B., Saunders, T. E., and Loose, M. (2020) Stochastic activation and bistability in a Rab GTPase regulatory network. Proc. Natl. Acad. Sci. U. S. A. 117, 6540–6549

25. Huotari, J., and Helenius, A. (2011) Endosome maturation. EMBO J. 30, 3481–500

26. Yorimitsu, T., and Klionsky, D. J. (2005) Autophagy: molecular machinery for self-eating. Cell Death Differ. 12 Suppl 2, 1542–52

27. Zimmermann, L., Stephens, A., Nam, S. Z., Rau, D., Kübler, J., Lozajic, M., Gabler, F., Söding, J., Lupas, A. N., and Alva, V. (2018) A Completely Reimplemented MPI Bioinformatics Toolkit with a New HHpred Server at its Core. J. Mol. Biol. 430, 2237–2243

28. Jumper, J., Evans, R., Pritzel, A., Green, T., Figurnov, M., Ronneberger, O., Tunyasuvunakool, K., Bates, R., Žídek, A., Potapenko, A., Bridgland, A., Meyer, C., Kohl, S. A. A., Ballard, A. J., Cowie, A., Romera-Paredes, B., Nikolov, S., Jain, R., Adler, J., Back, T., Petersen, S., Reiman, D., Clancy, E., Zielinski, M., Steinegger, M., Pacholska, M., Berghammer, T., Bodenstein, S., Silver, D., Vinyals, O., Senior, A. W., Kavukcuoglu, K., Kohli, P., and Hassabis, D. (2021) Highly accurate protein structure prediction with AlphaFold. Nature. 596, 583–589

29. Gautier, R., Douguet, D., Antonny, B., and Drin, G. (2008) HELIQUEST: a web server to screen sequences with specific-helical properties. Bioinformatics. 24, 2101–2102

30. Chorlay, A., and Thiam, A. R. (2020) Neutral lipids regulate amphipathic helix affinity for model lipid droplets. J. Cell Biol. 10.1083/jcb.201907099

31. Zinser, E., and Daum, G. (1995) Isolation and biochemical characterization of organelles from the yeast, Saccharomyces cerevisiae. Yeast. 11, 493–536

32. Schütter, M., Giavalisco, P., Brodesser, S., and Graef, M. (2020) Local Fatty Acid Channeling into Phospholipid Synthesis Drives Phagophore Expansion during Autophagy. Cell. 180, 135–149.e14

33. Li, L., Tong, M., Fu, Y., Chen, F., Zhang, S., Chen, H., Ma, X., Li, D., Liu, X., and Zhong, Q. (2021) Lipids and membrane-associated proteins in autophagy. Protein Cell. 12, 520–544

34. Klionsky, D. J., Abdel-Aziz, A. K., Abdelfatah, S., Abdellatif, M., Abdoli, A., Abel, S., Abeliovich, H., Abildgaard, M. H., Abudu, Y. P., Acevedo-Arozena, A., Adamopoulos, I. E., Adeli, K., Adolph, T. E., Adornetto, A., Aflaki, E., Agam, G., Agarwal, A., Aggarwal, B. B., Agnello, M., Agostinis, P., Tong, C.-K., et al. (2021) Guidelines for the use and interpretation of assays for monitoring autophagy (4th edition). Autophagy. 17, 1

35. Shibutani, S. T., and Yoshimori, T. (2014) A current perspective of autophagosome biogenesis. Cell Res. 24, 58–68

36. Bigay, J., and Antonny, B. (2012) Curvature, Lipid Packing, and Electrostatics of Membrane Organelles: Defining Cellular Territories in Determining Specificity. Dev. Cell. 23, 886–895

37. Joiner, A. M., Phillips, B. P., Yugandhar, K., Sanford, E. J., Smolka, M. B., Yu, H., Miller, E. A., and Fromme, J. C. (2021) Structural basis of TRAPPIII-mediated Rab1 activation. EMBO J. 40, e107607

38. Nguyen, N., Shteyn, V., and Melia, T. J. (2017) Sensing Membrane Curvature in Macroautophagy. J. Mol. Biol. 429, 457–472

39. Zens, B., Sawa-Makarska, J., and Martens, S. (2015) In vitro systems for Atg8 lipidation. Methods. 75, 37–43

40. Thomas, L. L., and Fromme, J. C. (2016) GTPase cross talk regulates TRAPPII activation of Rab11 homologues during vesicle biogenesis. J. Cell Biol. 215, 499–513

